# Fatigue influences the recruitment, but not structure, of muscle synergies

**DOI:** 10.1101/250522

**Authors:** P.A. Ortega-Auriol, T.F. Besier, W.D. Byblow, A.J.C. McMorland

**Author notes:** **Correspondence:** Pablo Ortega-Auriol.

## Abstract

The development of fatigue elicits multiple adaptations from the neuromuscular system. Muscle synergies are common patterns of neuromuscular activation that have been proposed as the building blocks of human movement. We wanted to identify possible adaptations of muscle synergies to the development of fatigue in the upper limb. Recent studies have reported that synergy structure remains invariant during the development of fatigue, but these studies did not examine isolated synergies. We propose a novel approach to characterize synergy adaptations to fatigue by taking advantage of the spatial tuning of synergies. This approach allows improved identification of changes to individual synergies that might otherwise be confounded by changing contributions of overlapping synergies. To analyse upper limb synergies we applied non-negative matrix factorization to 14 EMG signals from muscles of 11 participants performing isometric contractions. A preliminary multidirectional task was used to identify synergy directional tuning. A subsequent fatiguing task was designed to fatigue the participants in their synergies’ preferred directions. Both tasks provided virtual reality feedback of the applied force direction and magnitude, and were performed at 40% of each participant’s maximal voluntary force. Five epochs were analysed throughout the fatiguing task to identify progressive changes of EMG amplitude, median frequency, synergy structure, and activation coefficients. Three to four synergies were sufficient to account for the variability contained in the original data. Synergy structure was conserved with fatigue, but interestingly synergy activation coefficients decreased on average by 24.5% with fatigue development. EMG amplitude did not change systematically with fatigue, whereas EMG median frequency consistently decreased across all muscles. These results support the notion of a neuromuscular modular organization as the building blocks of human movement, with adaptations to synergy recruitment occurring with fatigue. When synergy tuning properties are considered, the reduction of activation of muscle synergies may be a reliable marker to identify fatigue.

## 1 Introduction

Fatigue has major implications for motor behaviour and task performance, with adaptations to fatigue occurring at central and peripheral levels of the neuromuscular system (Gandevia, 2001). Muscle synergies, which depend on covariations between levels of muscle activation, have been proposed as stable building blocks of human movement (d’Avella et al., 2003), but how their structure and recruitment are affected by adaptation to fatigue has yet to be determined.

One definition of fatigue is a decrease in muscle force that will lead to task failure (Enoka and Duchateau, 2008). The neuromuscular manifestations of fatigue reflect central and peripheral adaptations which can be jointly quantified by EMG during sustained muscle contractions (Bigland-Ritchie et al., 1979). Fatigue-related changes in myoelectric properties involve decreases of muscle conduction velocity (Enoka and Duchateau, 2008) and frequency of discharge of motor units (Dideriksen et al., 2011). In the EMG signal, fatigue reliably produces a decrease of the mean frequency (Bigland-Ritchie et al., 1981; Merletti et al., 1991) but has a variable effect on amplitude depending on a number of factors including the specific muscle involved and the level of contraction (Gerdle et al., 2000).

Changes of patterns of activation across muscles have been proposed as a central strategy to reduce the effects of fatigue (Enoka et al., 2011; Enoka and Duchateau, 2008). Four adaptations to muscle activation patterns have been described: activity alternation across synergistic muscles for low level contractions (< 5% MVC) (Kouzaki et al., 2002), co-activation with antagonists for moderate contractions (< 60% MVC) (Levenez et al., 2005), contralateral muscle activation (Todd et al., 2003), and increased variability of activation within the task parameters (Baudry et al., 2007). Motor control theory is currently lacking a unifying principle that explains these different adaptations.

There is a growing body of evidence that the central nervous system (CNS) controls the muscular system using a low-dimensional structure, composed of modules known as muscle synergies. A muscle synergy is a fixed pattern of co-activation of muscles that are driven by a common time-varying signal called the activation coefficient. Muscle synergies may be a strategy of the CNS to deal with the problem of motor redundancy (d’Avella et al., 2003). Muscle synergies have been found to be consistent across different natural movements in human and animal models (Bizzi et al., 2008a; Cheung et al., 2005).

The effect of fatigue on muscle synergies is not fully understood. For muscle synergies to be the building blocks of motor control, they must remain intact and well-defined across many different states of the motor system, including in the presence of fatigue. The aim of our study was to identify the adaptations of muscle synergies during the development of fatigue. To support the notion of synergies as building blocks of movement, we hypothesize that synergy structure is conserved with the development of fatigue. Consequently, to explain the differences in muscle activations during fatigue, adaptations should occur in the activation coefficients of muscle synergies. We studied adaptations to fatigue by comparing synergy structure and activation coefficients, as well as during the performance of fatiguing isometric upper limb contractions in humans.

## 2 Methods

### 2.1 Participants

We recruited eleven volunteer participants (**Table 1**); participants were young and healthy without any pathology that affected the upper limb, spine or posture. Volunteers were excluded if they reported neck, shoulder or arm pain (> 2 in a 1-10 verbal scale) within the last three months. The University of Auckland Human Participants Ethics Committee approved the research protocol and methods of the study (ref. 013218) and informed consent was gained prior to participation.

**Table 1:** Participant characteristics.

| ID | Age | Height (cm) | Weight (kg) | Arm Length (cm) | MVC force (N) |
| --- | --- | --- | --- | --- | --- |
| Mean | 25.8 | 170.9 | 67.4 | 65.8 | 74.9 |
| Median | 23.0 | 170.0 | 63.9 | 66.0 | 74.3 |
| SD | 4.7 | 7.9 | 12.9 | 4.6 | 22.1 |

### 2.2 Equipment

Forces generated by the participants were recorded at a handle instrumented with a 6-axis force-torque transducer (Omega160, ATI Industrial Automation, Apex, NC, USA) **(Figure 1.a)**. Force was sampled at 120 Hz using custom software based on Dragonfly acquisition system (Pittsburgh).

**Fig. 1:**
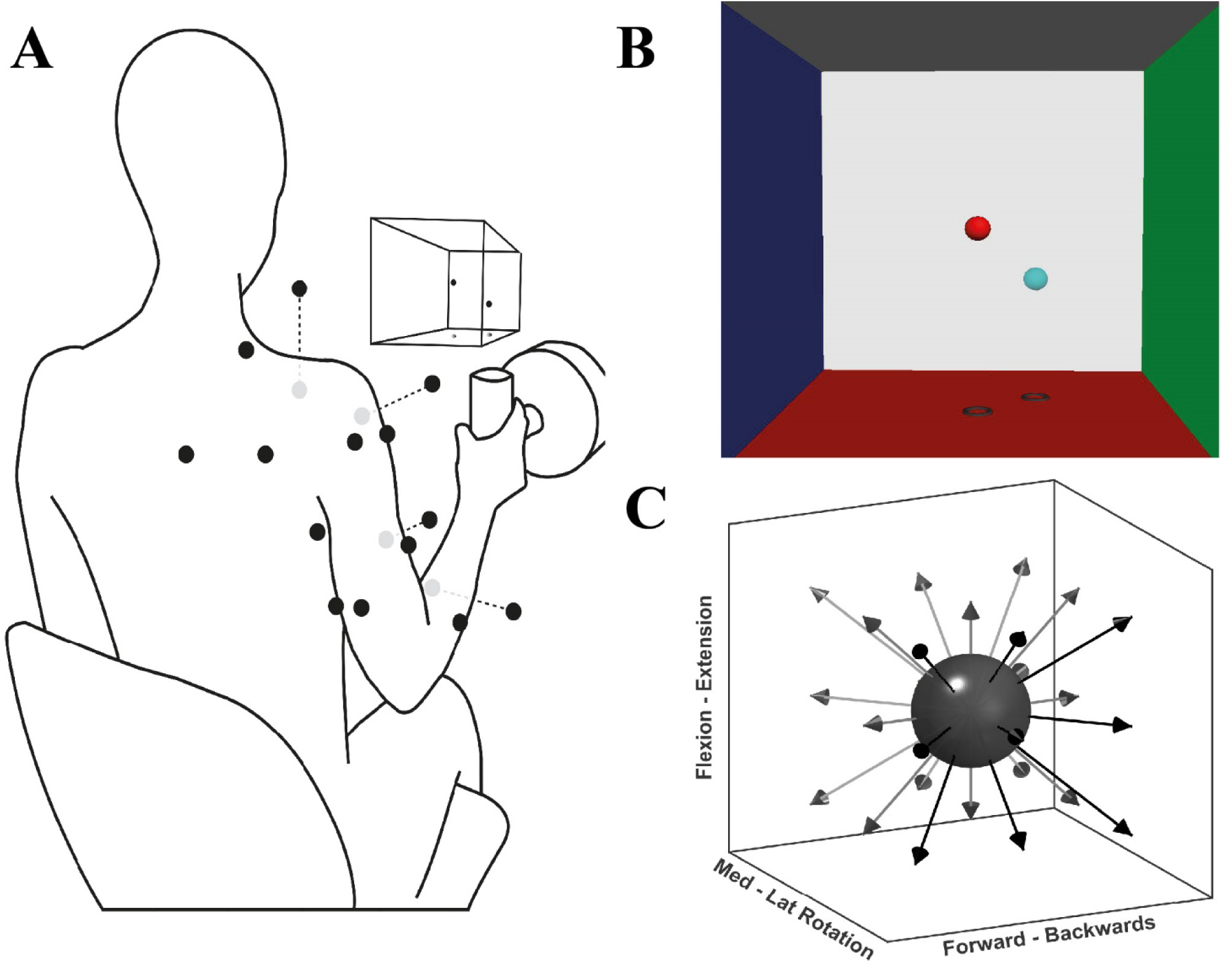
Experimental setup for the multidirectional and fatiguing tasks. (**A)**: Schematic illustration of EMG placement, VR feedback, and handle instrumented with a force transducer; black dots on the silhouette represents the EMG sensor placement, grey dots are sensors placed ventrally. (**B)** Screenshot of the VR feedback displayed on the screen in front of the participants, the centre of each VRF wall represents 100 N of applied force. (**C)** Schematic representation of the target directions during the multidirectional task.

Surface EMG signals were recorded with a wireless Trigno System (Delsys Inc., Boston, MA, USA). EMG activity was recorded from 14 major muscles of the shoulder, arm and forearm of the participant’s dominant upper limb: superior (ST) and middle trapezius (MT), subscapular (Sb), serratus anterior (SA), anterior (AD), middle (MD) and posterior deltoid (PD), pectoralis major (PM, clavicular fibres), short (BS) and long (BL) head of biceps brachii, long (TL) and lateral (TLat) heads of triceps brachii, extensor carpi radialis (WE), and flexor carpi radialis (WF). These muscles were chosen for the likely large contribution to tasks’ isometric contractions, as previously recommended to most accurately reconstruct synergies from a subset of muscles (Steele et al., 2013). Participants’ skin was prepared by rubbing a medical abrasive conductive paste (NuPrep, Weaver). Finally, electrode placement followed SENIAM and Cram’s recommendation guidelines (Criswell, 2010; Hermens et al., 1999). EMG signals were sampled at 2000 Hz, via a custom software interface.

### 2.3 Protocol

Participants performed three tasks: maximal voluntary force (MVF), multidirectional trials, and fatiguing trials. Each task consisted of isometric contractions of different time lengths and directions. During the tasks, participants were seated on a stool and grasped the instrumented handle that was positioned in front of their shoulder, at a distance of 40% of their respective arm length **(Figure 1.a)**. During the tasks, participants were instructed and encouraged to maintain their posture. To display the direction in which force had to be exerted, a custom virtual reality feedback (VRF) was developed **(Figure 1.b)**. The VRF consisted of two spheres in a 3D force space: the position of one sphere dynamically displaying the users applied force, and a fixed sphere serving as directional target and force level cue. The movement of the former was proportional to the resultant force exerted at the handle by the participant. The position of the target sphere was the desired vector direction with a distance from the origin equivalent to 40% of the MVF. Initially, participants were trained on the use of the force transducer-VRF interface by practising target matching in random directions and forces. Participants were then asked to perform shoulder external rotation producing MVF while seated with the upper arm next to the trunk, and the elbow at 90° degrees. Three MVF trials of 3 s were recorded, and the average MVF was determined from the peak forces.

During the multidirectional task, participants executed isometric trials in 26 directions evenly distributed around a sphere **(Figure 1.c)**. The goal of each trial was to match the movable sphere with the target one, applying a force of 40% of the MVF (+ 7 N.) for four seconds. If after three attempts of one minute, the participant was not able to achieve a continuous match of four seconds, the trial was considered a mistrial and excluded from further analysis. To prevent muscle fatigue a resting period of 20 s was given between trials.

For the fatiguing trials, participants were asked to perform one isometric contraction at 40% of the MVF until fatigue, for each of the significant synergies identified from the multidirectional task. The trials were performed in the preferred directions (PDs) of the extracted synergies. Synergy PDs are the directions for which a specific synergy shows the highest activation coefficient (see Data Analysis below). The CR10 Borg scale (Borg, 1998) was used to quantify the participant’s self-perceived exertion rate. The Borg scale is a subjective method to quantify fatigue development while performing a task. Participants reported their perceived exertion at the beginning of each trial and every one minute until fatigue was reached. Fatigue was reached when the force level dropped > 10 N for two consecutive seconds or when the participant declared it impossible to continue with the task, 10 on the Borg Scale. A rest period of 15 minutes was given between each fatiguing trial, allowing the muscles to return to an initial state.

### 2.4 Data Analysis

Data were first averaged and trimmed to obtain stable activation patterns from whole trials of the directional task and from epochs within the fatiguing task. With the fatigue data, fatigue parameters and synergies were calculated from each epoch. A significant number of synergies was identified and extracted from the concatenated data of the multidirectional trials, or independently from each epoch of the fatiguing trials. Code used for synergy analysis is available at https://github.com/ortegauriol/SynFAn.

#### 2.4.1 EMG processing

All data were analysed using custom scripts written in MATLAB 9.2 (MathWorks, Natick, MA, USA). EMG from the multidirectional trials were averaged for the intermediate two seconds of the target match period, obtaining a stable activation level. Fatiguing trials were trimmed and analysed in five epochs of 5% of the data, every 25% of the total trial time. Epochs were then rebinned into 100 data points. Signals were band-pass filtered (Butterworth, 2nd order, 5 - 400 Hz), demeaned, full-wave rectified, normalized by dividing all muscle activations by the maximum activation per trial (preserving the relative amplitude contribution of each muscle), converted to unit variance (Roh et al., 2012), and low-pass filtered again (Butterworth, 2nd order, 5 Hz) to obtain a signal envelope.

#### 2.4.2 EMG analysis

Non-negative matrix factorization (NMF) analysis (Lee and Seung, 1999) was used to extract synergies. In simple mathematical terms, NMF can be modelled as D = W*C, where D is the original data set, W the synergy structure or modes, and C the activation coefficient. For the analysis, NMF was implemented using the multiplicative rule (Berry et al., 2007), where each iteration gave a different W and C estimate, converging from the previous solution. The final solution was implemented as the result of 20 consecutive iterations with a difference of less than 0.01% among them (Roh et al., 2015).

Synergy analysis requires a pre-defined number of synergies. Therefore, to find an adequate solution, we iterated the number of synergies from one until 13 (number of muscles minus one). The concept behind muscle synergies is that a reduced dimensionality, or number of synergies in this case, is able to reconstruct complex higher dimensional behaviour. To determine a significant solution or number of synergies, the quality of the solution must be compared with the original data. We used the variance accounted for (VAF) metric to make this comparison (Cheung et al., 2005). VAF is defined in global (whole data set) and local (individual channels) scales. VAF was defined according to equation 1:

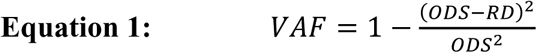

For global criteria, *ODS* represents the variance of the original data set, and *RD* the variance of the reconstructed data set. The local criterion of VAF is applied to each muscle independently: this involves replacing *ODS* with the variance of data from a single channel, and RD is replaced by the variance of the same reconstructed channel. The significant number of synergies was selected when global VAF => 90% and local VAF=>80%.

Once a significant number of synergies was determined, the PD of each synergy from the multidirectional task was calculated as the average of each trial’s direction vector scaled by the activation coefficient of that synergy during that trial (equation 2).

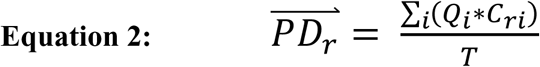

Where *Q*_*i*_ is the direction unit vector of the *i*th trial, *C*_*ri*_ is the activation coefficient of the *r*th synergy of the *i*th trial, and *T* is the total number of trials.

#### 2.4.3 Synergy clustering

To group similar synergies across participants in each epoch of the fatiguing task, synergies from all participants were pooled. Cluster analysis was applied to the pooled synergies using the K-medoids algorithm (Park and Jun, 2009), using the cosine function as the distance metric across clusters, and the Silhoutte index (Kaufman and Rousseeuw, 1990) to determine the correct number of clusters. Then, a mean synergy set per epoch was calculated by averaging each cluster. A second ‘sorting’ cluster analysis across epochs was applied to the calculated mean synergy sets to match similar mean synergies across epochs.

To test our first hypothesis that synergy structure is conserved with the development of fatigue, structure was compared across epochs by calculating the scalar product between synergies. Two synergies were defined as similar when the scalar product value was above the 95th percentile of a distribution of scalar products generated by comparing unstructured synergies. Given that synergy weights are constrained to have positive values between zero and one (normalized), there is a chance for spurious similarities. Using a threshold value from a by-chance distribution of scalar products (Roh et al. 2013) reduces the possibility of false positive similarity. To create shuffled unstructured synergies, we pooled all synergy structures from the fatiguing trials, and randomly shuffled weight values across epochs and muscles. Then, we calculated the scalar product between all shuffled synergies to obtain a baseline by-chance similarity distribution of scalar products.

#### 2.4.4 Fatigue parameters

EMG adaptations to fatigue were analysed for the muscles with the highest weight within each of the three synergy clusters: MT, AD, and ST. In theory, these muscles have a higher contribution to the performed contraction. This signal processing scheme allowed us to compare EMG amplitude, frequency, synergy structure, and activation coefficients along the development of fatigue. Fatigue parameters of power spectrum median frequency and signal amplitude of each epoch from the fatiguing trials were calculated by the Welch method (Welch, 1967) and RMS analysis respectively. All parameters were normalized to their value in the first epoch.

Separate one-way repeated measures ANOVAs were conducted with the dependent measures being the synergy activation coefficients and fatigue parameters (amplitude and median frequency) and the independent measure being the fatigue epoch. For each ANOVA, if significant differences were found, Bonferroni corrected paired t-tests were used to identify specific differences between epochs. Finally, to determine the trend of changes across epochs, a Pearson’s correlation analysis was used to describe the relationship between synergy coefficients and fatigue parameters from the muscles with the highest weight in each synergy. Statistics were performed using SPSS (version 24, IBM, NY).

## 3 Results

Representative raw and processed EMGs from a single subject during the multidirectional and fatiguing trials are displayed in **Figure 2.** All participants completed the multidirectional and fatiguing trials without missing any targets. For all participants, VAF analysis identified that three to four synergies (mean (SD) = 3.6 (0.5)) were sufficient to reconstruct the original muscle activation dataset from multidirectional trials.

**Fig. 2:**
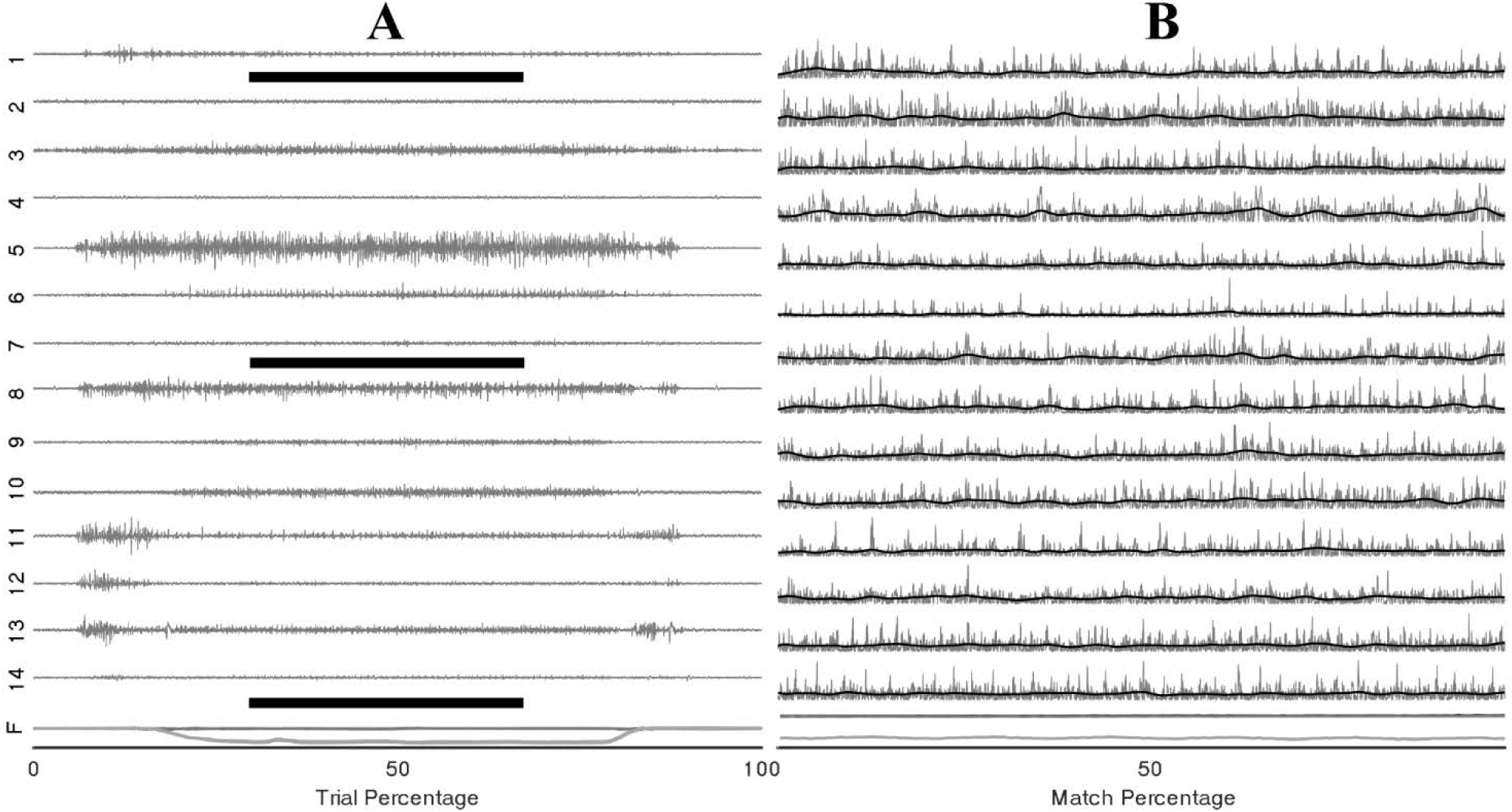
Example EMG and force traces from a single trial of the multidirectional task. (**A)**: Raw EMG signals of the 14 recorded muscles (1-14) and components of force vector (F) during a complete trial, black bars represent the period shown in (**B):** EMG and force traces trimmed to the central two seconds of the target-match period of the multidirectional task (grey); rectified and low-pass filtered signal (black).

### 3.1 Fatigue parameters

Changes of EMG median frequency, amplitude, and synergy activation coefficients were compared across epochs for each individual synergy. EMG median frequency (**Table 2**) changed in some but not all muscles with the development of fatigue. For MT median frequency: Maulchy’s test indicated that the assumption of sphericity was violated across the different epochs *χ*^2^ (9) = 86.4, p = 0.001, therefore Greenhouse-Geisser correction is reported (ε = 0.45). The results indicate that median frequency was different across epochs for MT (F[1.8, 70.6] = 3.92, p = 0.028, *ω*^2^ = 0.009). However, a post-hoc analysis did not find significance between the first and fifth epoch (highest mean difference 8%, Bonferroni-corrected paired t-test p = 0.15). For AD median frequency, again sphericity was violated (*χ*^2^ (9) = 85.9, p = 0.001) and a Greenhouse-Geisser correction of ε = 0.45 was used. Results suggest that there is no difference for the AD with the development of fatigue (F[1.8, 70.8] = 2.1, p = 0.13, *ω*^2^ = 0.005). Finally, for ST sphericity was violated [9] = 27.3, p = 0.001), with a Greenhouse-Geisser correction of ε = 0.75. ANOVA suggest that median frequency decreased with the development of fatigue for the ST (F [3, 116.5] = 28.264, p = 0.001, *ω*^2^ = 0.005). Post-hoc analysis found multiple differences (p < 0.05). These differences were between the 1^st^ and all other epochs (p = 0.001 for all), and between epochs 2-4, 2-5, and 4-5, (p = 0.001 for all). Fatigue development produced changes of parameters in some of the major contributors within synergies (**Figure 3**).

**Table 2:** Extracted fatigue parameters and synergy activation coefficients (mean (SD)) values. EMG amplitude and frequency is present as the average of the MT, AD, and ST muscles.

| Epoch | II | III | IV | V |
| --- | --- | --- | --- | --- |
| EMG amplitude | 100.6 (4.6) | 99.9 (7.5) | 97.3 (11.1) | 84.7 (14.8) |
| EMG median frequency | 95.7 (1.8) | 94.8(2.7) | 91.9(2.6) | 90.2 (2.9) |
| External rot. synergy | 97.1 (8.3) | 95.1 (5.5) | 94.1 (9.9) | 68.6 (10.1) |
| Flexion synergy | 91.9 (12.2) | 97.8 (11.7) | 82.7(18.7) | 80.5(13.2) |
| Extension synergy | 97.4 (11.3) | 92.8 (11.4) | 94.0 (10.3) | 77.4 (12.4) |

**Fig. 3:**
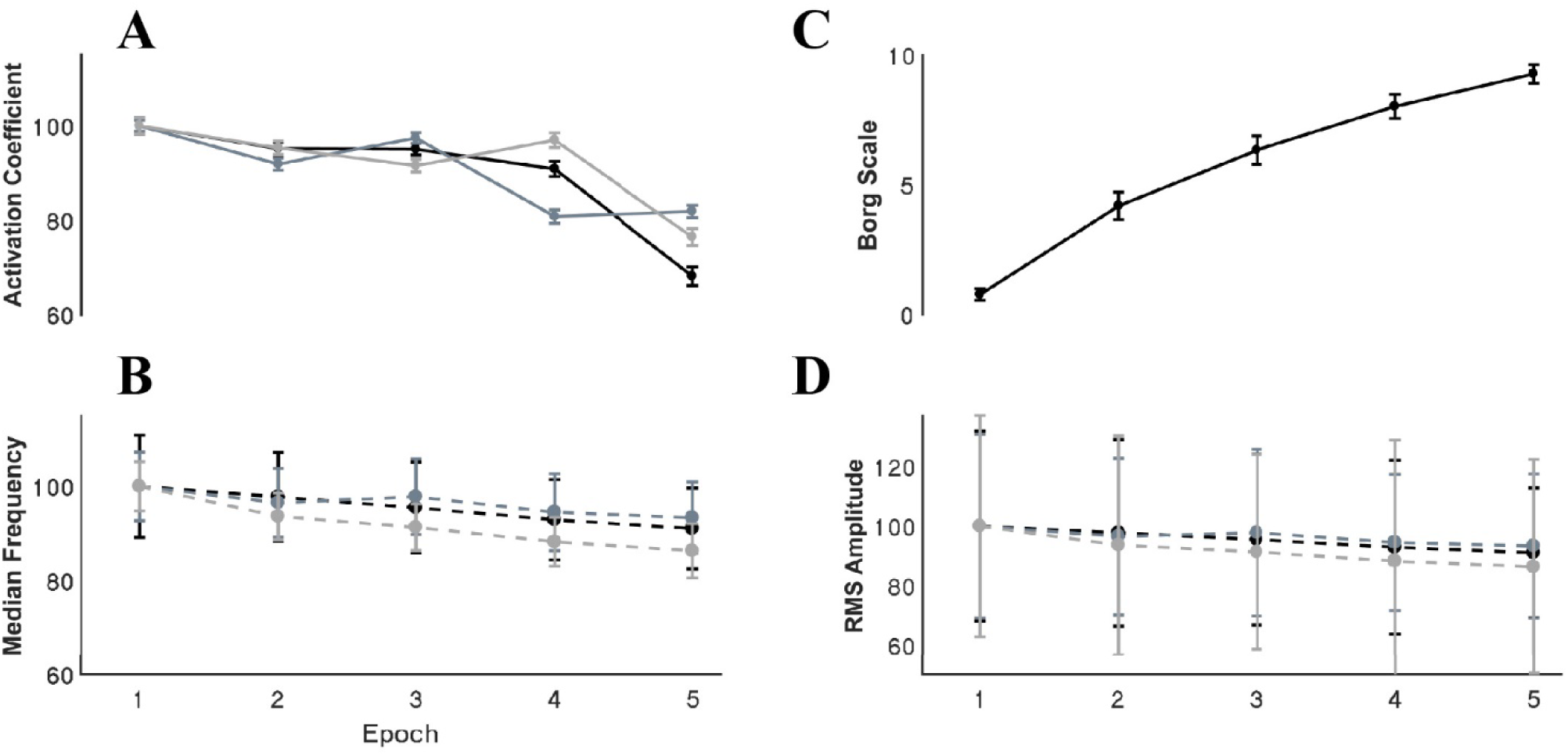
Activation coefficients and fatigue parameters through the development of fatigue. Error bars show the 95% confidence interval. For the activation coefficient panel, black, grey and light grey traces represent internal rotation, flexion and extension synergies respectively. For the EMG parameters the black, grey and light grey traces represents the MT, AD, and ST muscles respectively Signal RMS (**Table 2**) changed with the development of fatigue only for the MT muscle. Sphericity was violated (*χ*^2^[9] = 36.6, p = 0.001), consequently Greenhouse-Geisser correction was used (ε = 0.68). ANOVA reported a decrease of the amplitude in time with the development of fatigue (F[2.7, 105.9] = 10.2, p = 0.001, *ω*^2^ = 0.0015). Bonferroni-corrected t-test post-hoc analysis revealed several significant decreases of the Trapezius muscle with fatigue (epochs: 1-5 (p = 0.001), and 2-5 (p = 0.003) (**Figure 3**).

### 3.2 Synergies modulation

Synergies showed a distinctive tuning or modulation associated with direction of exertion: activation coefficients were highest for one particular direction, its preferred direction (PD), and decreased as the angle between the direction of exertion and the PD increased **(Figure 4a)**. After extracting synergies from the pooled fatiguing trials, cluster analysis identified four different clusters (**Figure 4b**). The PDs of these synergies were distributed approximately evenly throughout space. The predominant movement and muscle of these clusters were: external rotation (MT), flexion (AD), extension (ST), and internal rotation (PM).The second cluster analysis grouped synergies in three clusters per epoch, suggesting that synergies representing internal rotation became reclassified into the flexion and extension clusters.

**Fig. 4:**
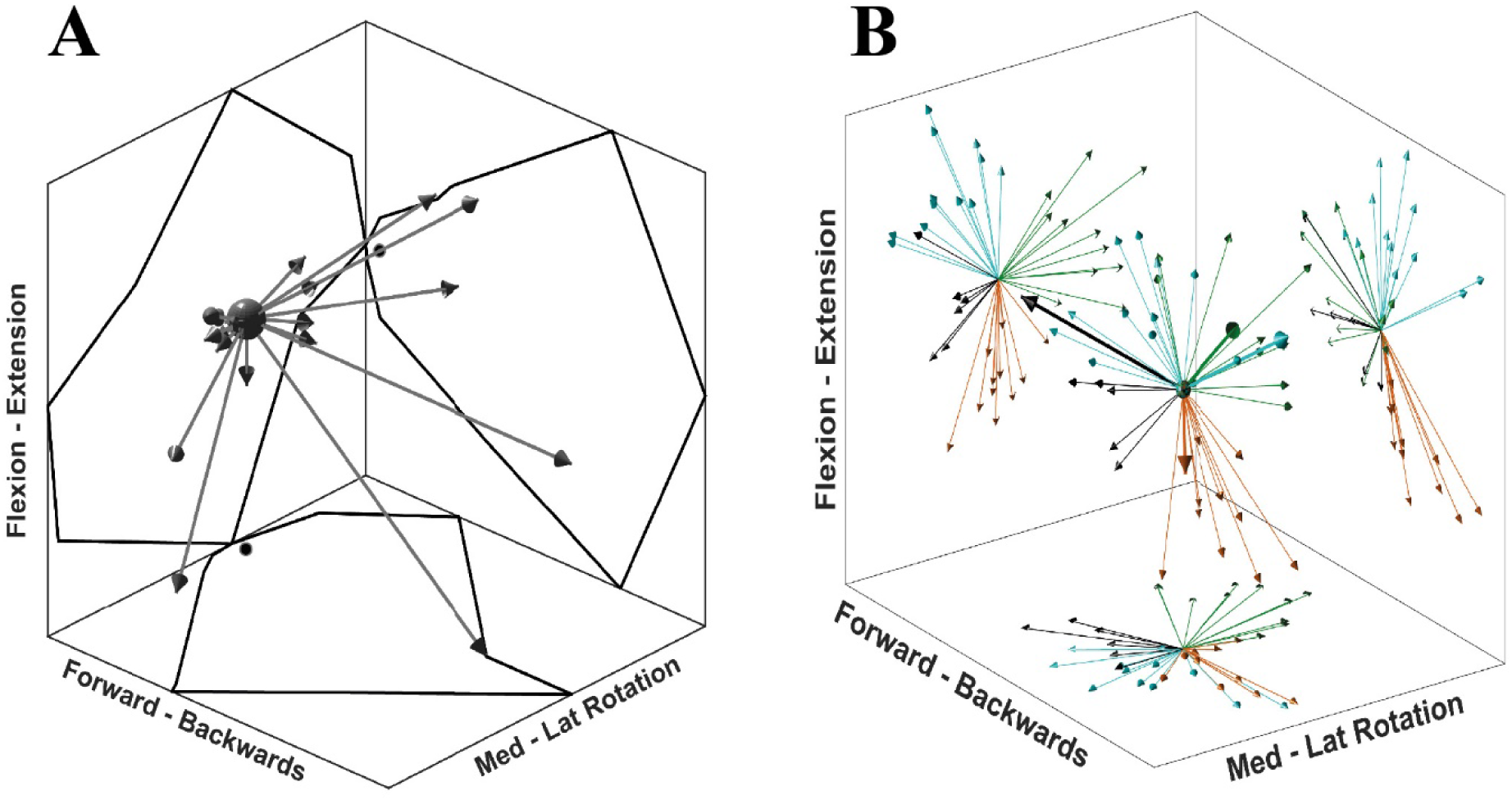
Synergy directional tuning and preferred directions. (**A)** A single synergy tuning over 26 directions of the multidirectional task. Each arrow represents a single target direction, and the length of the arrows is scaled to the synergy activation coefficient. The outline reflected on the graph walls displays the outer reach of the arrows, black dots on walls indicate the (0,0) coordinate. (**B)** 3D display of synergy PD for each synergy of all participants. Graph walls display a 2D projection of synergy directions showing the cluster distribution. Each colour represents a cluster identified from the first cluster analysis. Thicker arrows on the 3D display identify the synergies from a single, randomly-chosen participant.

### 3.3 Synergy structure

In agreement with our first hypothesis, synergy structure was conserved with the development of fatigue (**Figure 5**). Conservation of synergy structure was reflected by the high values of the scalar products between instances of an individual synergy across different epochs. Conservation of structure was analysed within each of the three mean synergy clusters derived from the sorting cluster analysis. Within each cluster, the mean scalar product was: S1 mean 0.98 (SD 0.009), S2 0.99 (0.007), and S3 0.97 (0.02). Finally, a paired t-test analysis revealed that the similarity within the synergy clusters was different from the similarity within a shuffled synergy group (T[59] = 23.6, p = 0.001, representing a large effect size *η^2^* = 0.90). This suggests that the similarity within synergy clusters is higher than would be expected by chance.

**Fig. 5:**
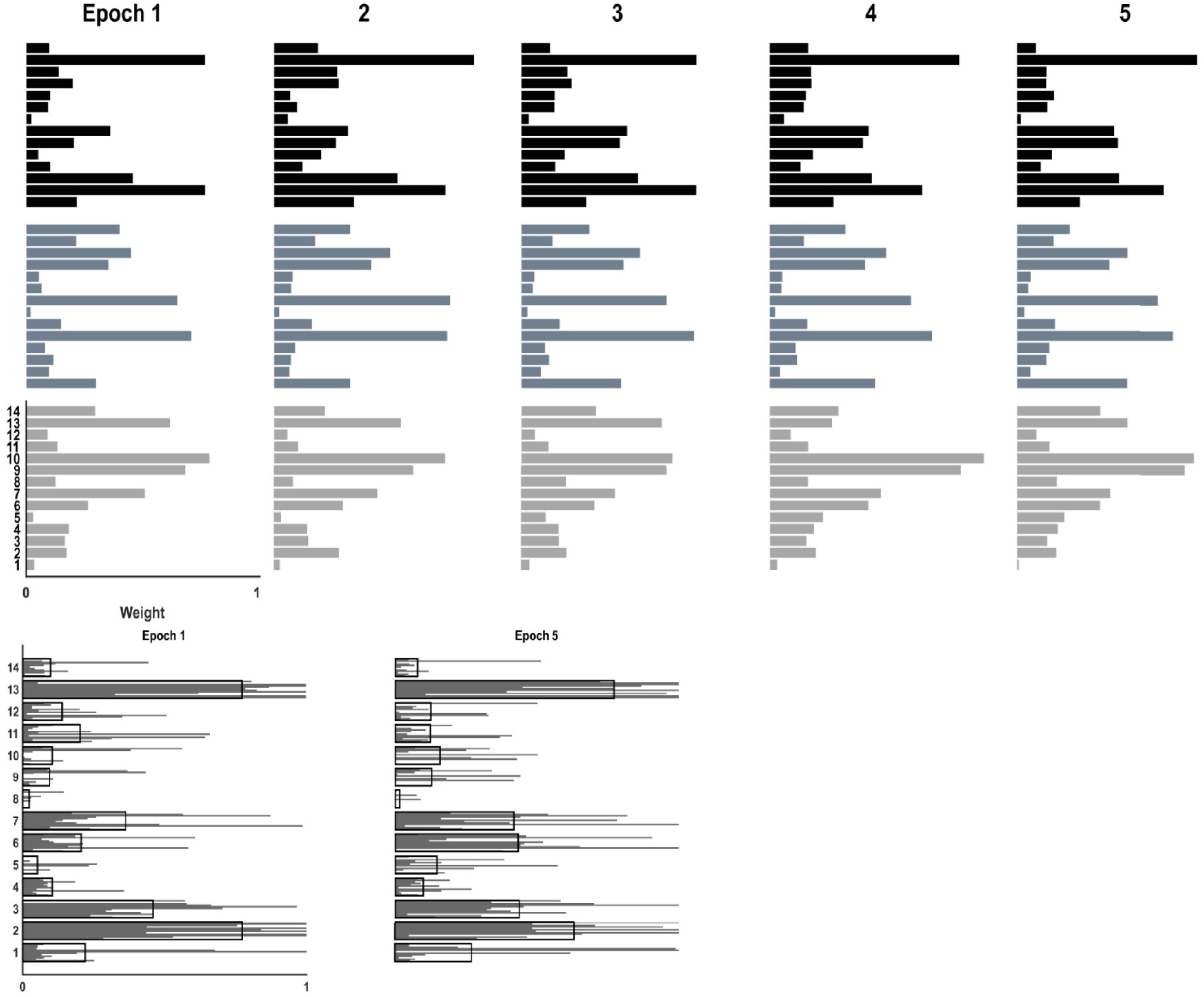
Synergy structure (weights) through the development of fatigue of the three identified synergies: internal rotation (black), flexion (grey), and extension (light grey). **(A)** The structure of the three synergies was conserved with the development of fatigue. **(B)** Variability of the structure of a single synergy, comparing the first and fifth epoch. Grey bars show the contribution of each muscle to the synergy identified in a single participant. Black-bordered bars indicate mean weights of the synergy cluster across all participants, which have been used for comparisons across epochs. The ordering of the subjects grey bars is the same for each muscle.

### 3.4 Synergy activation coefficients

Synergy activation coefficients decreased with the development of fatigue (**Table 2**, **Figure 4**). S1 analysis showed a conserved sphericity (χ^2^[9] = 13.6, p = 0.15). S1 activation coefficient showed a decrease with the development of fatigue (F[4, 40] = 4.2, p = 0.006, *ω*^2^ = 0.001). Post-hoc analysis revealed differences ranging between 3-31%; these were significant between epochs: 1-5 (p = 0.005), 2-5 (p = 0.05), 3-5 (p = 0.04). The mean S2 activation coefficient had a minor decrease of 18% of the normalized coefficient value between the first and fifth epoch (p = 0.22). Finally, S3 showed an even smaller decrease of 13% of its coefficient of activation.

To further explore the causes of the changes in activation coefficient while conserving synergy structure in terms of the relative changes to individual muscles, we examined the changes of EMG median frequency and amplitude in different muscles across epochs. We calculated the correlation coefficients of these parameters in two ways. Firstly, we compared the values from all muscles, and secondly, we focused on a muscle with the highest weight and another muscle randomly selected from those with an intermediate weight between 0.6 and 0.4. We did not examine a muscle with a low contribution to each synergy because we would expect the amplitude, and therefore signal-to-noise ratio, of their EMG signals to be low. When all the muscles were considered, the change of median frequency correlation was high (r = 0.95 (0.04)), suggesting that the median frequency of all muscles decreased with a similar trend regardless of the level of involvement of the muscle in the task. On the other hand, EMG amplitude correlation changes were moderate (r = 0.43 (0.4)). When looking at a high and medium contributor muscle to each synergy, the median frequency correlation average of the three synergies was high (r = 0.77 (0.08)), and signal amplitude correlation was low (r = 0.3 (0.5)).

Finally, to characterize the behaviour of synergy activation coefficients as a possible predictor of fatigue, correlation coefficients were calculated between synergy activation coefficients, fatigue parameters of the muscles, and Borg scale (**Figure 4**) with the highest weight of the correspondent synergy **(Table 3)**. Activation coefficients were correlated with mean frequency (all r > 0.7) and RPE (all < -0.7), while correlations with amplitude were inconsistent (r = 0.94 - 0.05).

**Table 3:** Correlation between synergy activation coefficients and other fatigue adaptation parameters.

| Synergy /Muscle | External rot. / MT | Flexion / AD | Extension / ST |
| --- | --- | --- | --- |
| Amplitude | 0.943 | 0.498 | 0.614 |
| Frequency | 0.838 | 0.957 | 0.708 |
| RPE | -0.771 | -0.829 | -0.715 |

#### 4 Discussion

Muscle synergies are proposed as the building blocks of human movement (Bizzi et al., 2008b). To fulfil this role, synergy structure should remain fixed, or at least known, by the intact central nervous system. The development of fatigue produces adaptations at central and peripheral levels of the neuromuscular system, which implies that synergy structure could change if counteracting adaptations did not exist. We hypothesize that synergy structure would be conserved with the development of fatigue, which would support the notion of synergies as fundamental units of motor control. It is important to note that fatigue induces adaptations in amplitude and frequency of the EMG signal, not only of the actuator muscles but also antagonists (Enoka and Duchateau, 2008). If synergies are purely the result of biomechanical constraints (Valero-Cuevas et al., 2009), changes in the signal amplitude across muscles would reflect changes in synergy structure.

In support of our hypothesis, we found that, with the development of fatigue demonstrated by the decrease of the median frequency and changes in amplitude of the signal, remarkably, synergy structure remains intact. However, synergy activation coefficients consistently decrease with the development of fatigue. This decrease correlates with characteristic adaptations found in the spectral analysis of the EMG signal, suggesting common mechanisms of regulation.

Adaptations to fatigue occur along the neuromuscular path from cortex to muscle, including changes at muscle, spinal, and cortical levels (Gandevia, 2001). During an isometric contraction, modulation of motoneuron activity is mediated by afferent inputs from peripheral receptors: muscle spindles (Macefield et al., 1993), Golgi tendon organs, and small diameter afferents (III and IV) (Hayward et al., 1991). Afferent input is partially responsible for the progressive decrease of the firing rate of motor units (Garland et al., 1988; Macefield et al., 1993; Vallbo, 1974) by inhibition, reducing facilitation, and presynaptic modulation (Gandevia, 2001). These peripheral afferents further decrease central drive that may itself be diminished with sustained contractions (Gandevia et al., 1996).

It is possible that synergies are generated and recruited at a central level with influence from the periphery. The outcome of these interacting adaptations to fatigue is a modification of muscle activation. In different muscles, we found that EMG amplitudes increased, decreased, or remained constant with fatigue. Remarkably, even in the presence of these uncorrelated changes of the EMG amplitude, synergy structure remained invariant. Similar behaviour is found in animal models; deafferentation results in drastic changes of EMG amplitude which can be explained by invariant synergy structure and only changing activation coefficients (Cheung et al., 2005). Our results support their notion of a feedback mechanism able to influence muscle synergy recruitment without changing the underlying structure. Synergy structure is also conserved across different natural movements of animals (Cheung et al., 2005) and humans (d’Avella et al., 2003). Therefore, the apparent universal conservation of synergy structure is consistent with this structure resulting from an underlying neuroanatomical circuitry at spinal or supra-spinal levels (Bizzi et al., 2008b), and the notion of hard, robustly encoded synergies from central sites.

To understand the intricate interactions that would influence synergy recruitment but conserve structure in the presence of fatigue, we need to consider the relationships between the known central and peripheral mechanisms of fatigue adaptation. Central fatigue is defined as the decrease of efferent drive from central sites, modulated by the influence of peripheral afferents (Gandevia, 2001). Experimentally, this can be demonstrated by an extra force output evoked by twitch interpolation (Allen et al., 1995; Bülow et al., 1995) or transcranial magnetic stimulation (Gandevia et al., 1996) that increases with fatigue. Similarly, synergy activation coefficients have been proposed as the reflection of central drive (Bizzi et al., 2008b; d’Avella et al., 2006). The decline of the synergy activation coefficients resembles that of central fatigue. Deafferentation of animal models produce changes of synergy recruitment (Cheung et al., 2005). Consequently, our findings of a decrease of synergy activation coefficients support the concept of an interaction among motor drive, spinal circuitry and afferent inputs (McCrea, 2001), integrating the proposed mechanisms of both fatigue and synergies. Afferent feedback influences the decrease of the muscle median frequency as an adaptation to fatigue and the strong correlation with synergy activation coefficients suggests a common modulatory mechanism.

The number of synergies, and more importantly, their structure, varies depending on the set of specific muscles included (Steele et al., 2013). To alleviate this effect it is important to include as many muscles as possible, in particular those that will contribute greatly to the task. Given the technical impossibility of acquiring all muscles of the upper limb, some structural differences from synergies that consider all muscles might underlie our results. Nevertheless, we included 14 muscles, mainly of the shoulder region, considering their probable muscle contribution to isometric contractions. Brachioradialis is one deeper muscle that would potentially make a significant contribution to synergy structure that is not easily recorded from using surface EMG. Another concern with the estimation of synergies is that the extracted number of synergies depends critically on the complexity of the task. Dynamic contractions utilize a higher number of synergies (Cheung et al., 2005, 2009; d’Avella et al., 2006) than less complex behaviour like isometric contractions. The amount of explored space is also relevant for the number of synergies, and it seems that this effect can outweigh the increase in number of synergies arising from more complex movement: from two synergies in dynamic exploration of a single plane (Muceli et al., 2010) to six synergies for movements in multiple planes and directions (Cheung et al., 2009). We found that three to four synergies were able to reconstruct the original EMG sets from the multidirectional trials. This number is similar to other studies (Roh et al., 2012, 2013; Steele et al., 2013) that have performed isometric contractions with two to four times the number of directions considered in our multidirectional task. Our task was more constrained in terms of force directions because our aim was not to find all available synergies. Nevertheless, we did likely obtain most synergies involved in performing isometric contractions with the upper limb.

Synergies seem to be a more reliable measure of fatigue development, highly correlated with parameters that have shown homogeneous changes with fatigue development. Synergy analysis implies the search for spatiotemporal patterns of muscle activity (McMorland et al., 2015), considering signal amplitude a key element of analysis. Our results show mixed changes of the EMG signal amplitudes within a specific synergy, corroborated by a moderate correlation between amplitude changes. Within EMG adaptations to fatigue, signal amplitude behaves inconsistently (De Luca, 1984; Dimitrova and Dimitrov, 2003; Gerdle et al., 2000), therefore it remains an unreliable measure of fatigue (Hultman and Sjöholm, 1983; Vøllestad, 1997). It is relevant to consider that a decline in force with fatigue does not directly imply a decline of the EMG amplitude (Merton, 1954). Thus, synergy activation coefficients seem like a better alternative to EMG amplitude although they require a slightly more elaborate analysis. In contrast, EMG spectral median frequency shows a consistent decrease with the development of fatigue (Bigland-Ritchie et al., 1981). Our results show an equivalent behaviour across all muscles, with a high correlation within a single synergy. However, this decrease shows certain variability across muscles and, once normalized, is not as great as the decrease of the synergy activation coefficients. EMG power spectrum is a compound characteristic affected by the intracellular action potential, motor unit potential, and consequently from motor drive and efferent signals (Dimitrova and Dimitrov, 2003). Synergy analysis looks at the modular control of fixed activation patterns of many muscles, thus it is likely that fewer relevant variables affect its behaviour. Although synergy analysis does not directly consider the EMG power spectrum, the correlation between EMG median frequency and synergy activation coefficients may reflect shared regulatory mechanisms.

Our experimental setup presents several advantages over other attempts to characterize synergy adaptations to fatigue (Smale et al., 2016; Turpin et al., 2011). First, by performing the fatiguing trials in the directions of synergy PDs, we have isolated the effects of fatigue to a single synergy at a time. This approach restricts confounding effects associated with load sharing across synergies, and maximizes the extent to which fatigue influences only the components relating to activation of the single synergy in question. Unspecific fatigue development of synergies, as seen previously in studies examining functional movements, may lead to wrong interpretations of outcomes and the underlying mechanisms. Secondly, we analysed adaptations to fatigue across multiple epochs, improving the opportunity to quantify changes throughout the development of fatigue. We see this as an improvement of the analysis of global measures during fatigue development. Analysis only considering whole trials might impede the identification of changes to synergy structure and activation coefficients. Turpin and colleagues (Turpin et al., 2011), using such an approach, have previously found that there were no changes in either the structure or activation coefficients during a fatiguing task. These two main advantages allowed us to analyse the effects of fatigue on synergy structure and activation coefficients in time, providing a better perspective of synergy adaptations to fatigue.

## 5 Conclusion

The invariability of synergy structure supports the notion of synergies as a robust mechanism of motor control. Our study demonstrates a novel approach able to detect the adaptations of synergies to fatigue by identifying decreases of synergy activation coefficients. Synergies’ adaptations to fatigue seem to be mediated by common neuromuscular regulatory mechanisms coordinating at central and peripheral levels. If synergy tuning is considered, synergy activation coefficients are likely to be a valuable approach to assess fatigue under isometric conditions.

## 6 Acknowledgments

## 7 Author contribution statement

PO-A and AM contributed to the original concept, design and technical development of the work. PO-A acquired and analysed the data, and wrote the draft of the work. All authors contributed with the discussion development. AM, WB and TB contributed with improvement to analysis and data collection, and revised the draft. AM and PO-A constructed the final version of the manuscript.

## References

Allen, G. M., Gandevia, S. C., and McKenzie, D. K. (1995). Reliability of measurements of muscle strength and voluntary activation using twitch interpolation. Muscle Nerve 18, 593–600.

Baudry, S., Klass, M., Pasquet, B., and Duchateau, J. (2007). Age-related fatigability of the ankle dorsiflexor muscles during concentric and eccentric contractions. Eur. J. Appl. Physiol. 100, 515–525.

Berry, M. W., Browne, M., Langville, A. N., Pauca, V. P., and Plemmons, R. J. (2007). Algorithms and applications for approximate nonnegative matrix factorization. Comput. Stat. Data Anal. 52, 155–173. doi:10.1016/j.csda.2006.11.006.

Bigland-Ritchie, B., Donovan, E. F., and Roussos, C. S. (1981). Conduction velocity and EMG power spectrum changes in fatigue of sustained maximal efforts. J. Appl. Physiol. 51, 1300–1305.

Bigland-Ritchie, B., Jones, D. A., and Woods, J. J. (1979). Excitation frequency and muscle fatigue: Electrical responses during human voluntary and stimulated contractions. Exp. Neurol. 64, 414–427. doi:10.1016/0014-4886(79)90280-2.

Bizzi, E., Cheung, V. C. K., d’Avella, A., Saltiel, P., and Tresch, M. (2008a). Combining modules for movement. Brain Res. Rev. 57, 125–133. doi:10.1016/j.brainresrev.2007.08.004.

Bizzi, E., Cheung, V. C. K., d’Avella, A., Saltiel, P., and Tresch, M. (2008b). Combining modules for movement. Brain Res. Rev. 57, 125–133. doi:10.1016/j.brainresrev.2007.08.004.

Borg, G. (1998). Borg’s Perceived Exertion and Pain Scales. Champaign, IL: Human Kinetics.

Bülow, P. M., Jørregaard, J., Mehlsen, J., and Danneskiold-Samsøe, B. (1995). The twitch interpolation technique for study of fatigue of human quadriceps muscle. J. Neurosci. Methods 62, 103–109. doi:10.1016/0165-0270(95)00062-3.

Cheung, V. C. K., d’Avella, A., Tresch, M. C., and Bizzi, E. (2005). Central and sensory contributions to the activation and organization of muscle synergies during natural motor behaviors. J. Neurosci. Off. J. Soc. Neurosci. 25, 6419–6434. doi:10.1523/JNEUROSCI.4904-04.2005.

Criswell, E. (2010). Cram’s introduction to surface electromyography. Jones & Bartlett Publishers.

d’Avella, A., Portone, A., Fernandez, L., and Lacquaniti, F. (2006). Control of fast-reaching movements by muscle synergy combinations. J. Neurosci. Off. J. Soc. Neurosci. 26, 7791–7810. doi:10.1523/JNEUROSCI.0830-06.2006.

d’Avella, A., Saltiel, P., and Bizzi, E. (2003). Combinations of muscle synergies in the construction of a natural motor behavior. Nat. Neurosci. 6, 300–308. doi:10.1038/nn1010.

De Luca, C. J. (1984). Myoelectrical manifestations of localized muscular fatigue in humans. Crit. Rev. Biomed. Eng. 11, 251–279.

Dideriksen, J. L., Enoka, R. M., and Farina, D. (2011). Neuromuscular adjustments that constrain submaximal EMG amplitude at task failure of sustained isometric contractions. J. Appl. Physiol. 111, 485–494. doi:10.1152/japplphysiol.00186.2011.

Dimitrova, N. A., and Dimitrov, G. V. (2003). Interpretation of EMG changes with fatigue: facts, pitfalls, and fallacies. J. Electromyogr. Kinesiol. 13, 13–36. doi:10.1016/S1050-6411(02)00083-4.

Enoka, R. M., Baudry, S., Rudroff, T., Farina, D., Klass, M., and Duchateau, J. (2011). Unraveling the neurophysiology of muscle fatigue. J. Electromyogr. Kinesiol. 21, 208–219. doi:10.1016/j.jelekin.2010.10.006.

Enoka, R. M., and Duchateau, J. (2008). Muscle fatigue: what, why and how it influences muscle function. J. Physiol. 586, 11–23.

Gandevia, S. C. (2001). Spinal and Supraspinal Factors in Human Muscle Fatigue. Physiol. Rev. 81, 1725–1789.

Gandevia, S. C., Allen, G. M., Butler, J. E., and Taylor, J. L. (1996). Supraspinal factors in human muscle fatigue: evidence for suboptimal output from the motor cortex. J. Physiol. 490, 529–536.

Garland, S. J., Garner, S. H., and McComas, A. J. (1988). Reduced voluntary electromyographic activity after fatiguing stimulation of human muscle. J. Physiol. 401, 547–556. doi:10.1113/jphysiol.1988.sp017178.

Gerdle, B., Larsson, B., and Karlsson, S. (2000). Criterion validation of surface EMG variables as fatigue indicators using peak torque: a study of repetitive maximum isokinetic knee extensions. J. Electromyogr. Kinesiol. 10, 225–232. doi:10.1016/S1050-6411(00)00011-0.

Hayward, L., Wesselmann, U., and Rymer, W. Z. (1991). Effects of muscle fatigue on mechanically sensitive afferents of slow conduction velocity in the cat triceps surae. J. Neurophysiol. 65, 360–370.

Hermens, H. J., Freriks, B., Merletti, R., Stegeman, D. F., Blok, J., Rau, G., et al. (1999). European recommendations for surface ElectroMyoGraphy : results of the SENIAM project. Enschede: Roessingh Research and Development.

Hultman, E., and Sjöholm, H. (1983). Electromyogram, force and relaxation time during and after continuous electrical stimulation of human skeletal muscle in situ. J. Physiol. 339, 33–40. doi:10.1113/jphysiol.1983.sp014700.

Kaufman, L., and Rousseeuw, P. (1990). Finding Groups in Data: An Introduction to Cluster Analysis. Wiley-Interscience Available at: http://www.amazon.ca/exec/obidos/redirect?tag=citeulike09-20&path=ASIN/0471735787 [Accessed November 23, 2017].

Kouzaki, M., Shinohara, M., Masani, K., Kanehisa, H., and Fukunaga, T. (2002). Alternate muscle activity observed between knee extensor synergists during low-level sustained contractions. J. Appl. Physiol. 93, 675–684.

Lee, D. D., and Seung, H. S. (1999). Learning the parts of objects by non-negative matrix factorization. Nature 401, 788–791. doi:10.1038/44565.

Levenez, M., Kotzamanidis, C., Carpentier, A., and Duchateau, J. (2005). Spinal reflexes and coactivation of ankle muscles during a submaximal fatiguing contraction. J. Appl. Physiol. 99, 1182–1188.

Macefield, V. G., Gandevia, S. C., Bigland-Ritchie, B., Gorman, R. B., and Burke, D. (1993). The firing rates of human motoneurones voluntarily activated in the absence of muscle afferent feedback. J. Physiol. 471, 429–443.

McCrea, D. A. (2001). Spinal circuitry of sensorimotor control of locomotion. J. Physiol. 533, 41–50. doi:10.1111/j.1469-7793.2001.0041b.x.

McMorland, A. J. C., Runnalls, K. D., and Byblow, W. D. (2015). A Neuroanatomical Framework for Upper Limb Synergies after Stroke. Front. Hum. Neurosci. 9. doi:10.3389/fnhum.2015.00082.

Merletti, R., Lo Conte, L. R., and Orizio, C. (1991). Indices of muscle fatigue. J. Electromyogr. Kinesiol. 1, 20–33. doi:10.1016/1050-6411(91)90023-X.

Merton, P. A. (1954). Voluntary strength and fatigue. J. Physiol. 123, 553–564. doi:10.1113/jphysiol.1954.sp005070.

Park, H.-S., and Jun, C.-H. (2009). A simple and fast algorithm for K-medoids clustering. Expert Syst. Appl. 36, 3336–3341. doi:10.1016/j.eswa.2008.01.039.

Pittsburgh, U. Dragonfly Messaging - The fastest way to connect ad-hoc software modules. Available at: http://dragonfly-msg.org [Accessed October 25, 2017].

Roh, J., Rymer, W. Z., and Beer, R. F. (2012). Robustness of muscle synergies underlying three-dimensional force generation at the hand in healthy humans. J. Neurophysiol. 107, 2123–2142. doi:10.1152/jn.00173.2011.

Roh, J., Rymer, W. Z., and Beer, R. F. (2015). Evidence for altered upper extremity muscle synergies in chronic stroke survivors with mild and moderate impairment. Front. Hum. Neurosci. 9. doi:10.3389/fnhum.2015.00006.

Smale, K. B., Shourijeh, M. S., and Benoit, D. L. (2016). Use of muscle synergies and wavelet transforms to identify fatigue during squatting. J. Electromyogr. Kinesiol. 28, 158–166. doi:10.1016/j.jelekin.2016.04.008.

Steele, K. M., Tresch, M. C., and Perreault, E. J. (2013). The number and choice of muscles impact the results of muscle synergy analyses. Front. Comput. Neurosci. 7. doi:10.3389/fncom.2013.00105.

Todd, G., Petersen, N. T., Taylor, J. L., and Gandevia, S. (2003). The effect of a contralateral contraction on maximal voluntary activation and central fatigue in elbow flexor muscles. Exp. Brain Res. 150, 308–313.

Turpin, N. A., Guével, A., Durand, S., and Hug, F. (2011). Fatigue-related adaptations in muscle coordination during a cyclic exercise in humans. J. Exp. Biol. 214, 3305–3314. doi:10.1242/jeb.057133.

Valero-Cuevas, F. J., Venkadesan, M., and Todorov, E. (2009). Structured variability of muscle activations supports the minimal intervention principle of motor control. J. Neurophysiol. 102, 59–68. doi:10.1152/jn.90324.2008.

Vallbo, A. B. (1974). Afferent Discharge from Human Muscle Spindles in Non-Contracting Muscles. Steady State Impulse Frequency as a Function of Joint Angle. Acta Physiol. Scand. 90, 303–318. doi:10.1111/j.1748-1716.1974.tb05593.x.

Vøllestad, N. K. (1997). Measurement of human muscle fatigue. J. Neurosci. Methods 74, 219–227.

Welch, P. D. (1967). The use of fast Fourier transform for the estimation of power spectra: A method based on time averaging over short, modified periodograms. IEEE Trans. Audio Electroacoustics 15, 70–73. doi:10.1109/TAU.1967.1161901.

